# Contrasting Variable and Stable Subsurface Microbial Populations: an ecological time series analysis from the Deep Mine Microbial Observatory, South Dakota, USA

**DOI:** 10.1101/2020.09.15.298141

**Authors:** Magdalena R. Osburn, Caitlin P. Casar, Brittany Kruger, Lily Momper, Theodore M. Flynn, Jan P. Amend

## Abstract

The deep subsurface contains a vast reservoir of microbial life. While recent studies have revealed critical details about this biosphere including the sheer diversity of microbial taxa and their metabolic potential, long-term monitoring of deep subsurface microbial populations is rare, thus limiting our understanding of subsurface microbial population dynamics. Here we present a four-year time series analysis of subsurface microbial life from the Deep Mine Microbial Observatory (DeMMO), Lead, SD, USA. We find distinct and diverse populations inhabiting each of 6 sites over this ~1.5 km deep slice of terrestrial crust, corresponding to distinct geochemical habitats. Alpha diversity decreases with depth and beta diversity measures clearly differentiate samples by site over time, even during substantial perturbations. Population dynamics are driven by a subset of variable (and often relatively abundant) OTUs, but the vast majority of detected OTUs are stable through time, constituting a core microbial community. The phylogenetic affiliations of both stable and variable taxa, including putative sulfate reducers, methanogens, spore formers, and many uncultivated lineages, are similar to those found previously in subsurface environments. This work reveals the dynamic nature of the terrestrial subsurface, contributing to a more holistic understanding than can be achieved when viewing shorter timeframes.

**Originality-Significance Statement:** This four-year record of deep mine microbial diversity and geochemistry is the first of its kind and allows for direct investigation of temporal trends in deep subsurface biogeochemistry. We identify disparate populations of variable and stable taxa, suggesting the presence of a core deep subsurface microbiome with unique niche partitioning.

## 1. Introduction

The deep subsurface hosts a large and diverse microbial community. Pioneering work in mines, boreholes, and springs has painted a picture of the deep surface as a habitable landscape rather than vast expanses of sterile rock (see reviews by (Pedersen, 2000; Fredrickson and Onstott, 2001; Amend and Teske, 2005; Fredrickson and Balkwill, 2006; Colwell and D’Hondt, 2013)). Estimates of the biomass of this realm have been continually refined with new data and now suggest that it is one of the largest ecosystems on the planet, constituting 15-23 Pg of carbon (Whitman *et al*., 1998; McMahon and Parnell, 2013; Magnabosco *et al*., 2018; Bar-On *et al*., 2018; Flemming and Wuertz, 2019). The deep terrestrial (or continental) biosphere is structurally complex owing to heterogenous geological environments produced via tectonic processes. Here, the environments and geochemical gradients available as microbial habitats and energy sources are shaped by hydrological connectivity in aquifers and fracture-based mixing. However, large open questions remain regarding the stability and connectivity of these intraterrestrial microbial communities.

A key technological advance toward understanding the microbial diversity of the deep terrestrial subsurface has been the application of DNA sequencing technologies to planktonic and attached biomass. Early efforts in this sphere revealed strong gradients in microbial taxa with depth (Pedersen, 2001; Haveman and Pedersen, 2002), unusual monophyletic communities (Chivian *et al*., 2008), and stark differences between attached and planktonic biomass (Pedersen, 1996; Lehman *et al*., 2001). More recent applications of second and third generation sequencing technologies have expanded on these results, revealing vast diversity of subsurface taxa particularly in uncultivated groups and spatial separation of populations with depth and between aquifers (Anantharaman *et al*., 2016; Jungbluth *et al*., 2016; Probst *et al*., 2017; Momper, Jungbluth, Lee, and Amend, 2017a; Probst *et al*., 2018), as well as key as genomic adaptations in deep intraterrestrials (Lau *et al*., 2016; Momper *et al*., 2018).

Long term datasets pertaining to the deep subsurface are rare, with studies generally consisting of either individual time points or sampling over a single field campaign. Probst et al. (2018) sampled fluid from Crystal Geyser over the course of a five-day eruptive cycle (Probst *et al*., 2017). They found distinct microbial assemblages sourcing from different aquifers (from a maximum of 800 m) over the course of the eruption including many uncultivated archaea and bacteria. Benk et al. (2019) targeted the very shallow subsurface over a three-year time series in concert with DOM analysis and found considerable temporal variability in the alpha diversity of groundwater communities that was correlated with the alpha diversity of DOM itself (Benk *et al*., 2019). Kadnikov et al. (2018) accessed a Siberian borehole over a five-year time period, but do not report a time series analysis of microbial diversity (Kadnikov *et al*., 2018).

The Sanford Underground Research Facility (SURF) hosted in the former Homestake Gold Mine, Lead SD is a key portal into the deep terrestrial subsurface. Accessible levels extend from 91 m to 1.5 km below surface with flooded levels down to 2.5 km. Previous sequencing efforts at SURF have focused on mine soils (Rastogi *et al*., 2009; 2010) and industrially relevant metabolisms (Rastogi *et al*., 2013). A detailed comparison between the geochemistry and microbial diversity of planktonic fluids was published in (Osburn *et al*., 2014), followed by a comparison between fluid and rock communities (Momper, Kiel Reese, *et al*., 2017), and a metagenomic analysis of two borehole sites (Momper, Jungbluth, Lee, and Amend, 2017a). A recent study of newly drilled boreholes finds distinct microbial communities in extremely close proximity and uses microbial similarly to identify fracture connectivity (Y. Zhang *et al*., 2019). These studies illustrate the vast diversity of microbes across different SURF locations and their metabolic potential, but lack a temporal element.

The Deep Mine Microbial Observatory (DeMMO) was established by the NASA Astrobiology Institute *Life Underground* team to facilitate long term sampling of the deep biosphere (Osburn *et al*., 2019). Six boreholes (D1-D6) are situated on different levels, at 244 m D1 & D2, 610 m D3, 1250 m D4, and 1478 m D5 & D6, each tapping fluid with distinct geochemistry, following the broad hydrological windows modeled in Murdoch et al. (2011). Osburn et al. (2019) describes the chemistry of these sites over a two-year period and find that fluids from each site are distinct and stable through time. Chemical similarities between the fluids relate to level of water-rock interaction grouping sites D1 and D3 based on longer residence times compared to D2, D4, and D5. D6 is distinct with signatures of long-term isolation (high conductivity, Na^+^, SO4^2-^, Fe^2+^, reduced gases). D4 and D5 are chemically similar (high NO_3^-^_, NH4^+^, H2S, in Na^+^ dominated fluid) and 3D maps of borehole trajectories reveal that these holes tap a proximal fluid source. Here we describe and evaluate a time series of 16S rRNA-derived microbial community composition and corresponding geochemistry collected over the course of four years during and years after the installation of the DeMMO network.

## 2. Results

### 2.1 Microbial diversity of DeMMO sites

High throughput 16S rRNA sequencing of DeMMO borehole fluids reveal diverse microbial communities that are distinct by site, but broadly consistent through time (Fig. 1). Each site is dominated by bacteria, with archaea occurring in lower relative abundance. Particularly abundant are sequences that could not be classified (*see experimental procedures*), as well as uncultured phyla and members of the Candidate Phyla Radiation (CPR, also known as the Patescibacteria). The taxa comprising more than 10% of each site are as follows: D1, unassigned OTUs (24%), Betaproteobacteria (22%), and Omnitrophicaeota (20%); D2, Betaproteobacteria (27%), unassigned OTUs (15%), and Deltaproteobacteria (10%); D3, Beta- (17%) and Deltaproteobacteria (12%), Nitrospirae (11%), unassigned OTUs (14%); D4, Betaproteobacteria (27%), Omnitrophicaeota (16%), and Gammaproteobacteria (10%); D5, Beta- (19%) and Deltaproteobacteria (15%); D6, Deltaproteobacteria (33%), Firmicutes (28%), and Bacteroidetes (10%). The numerous taxa which occur at lower abundance and changes in their distribution with space and time will be discussed below.

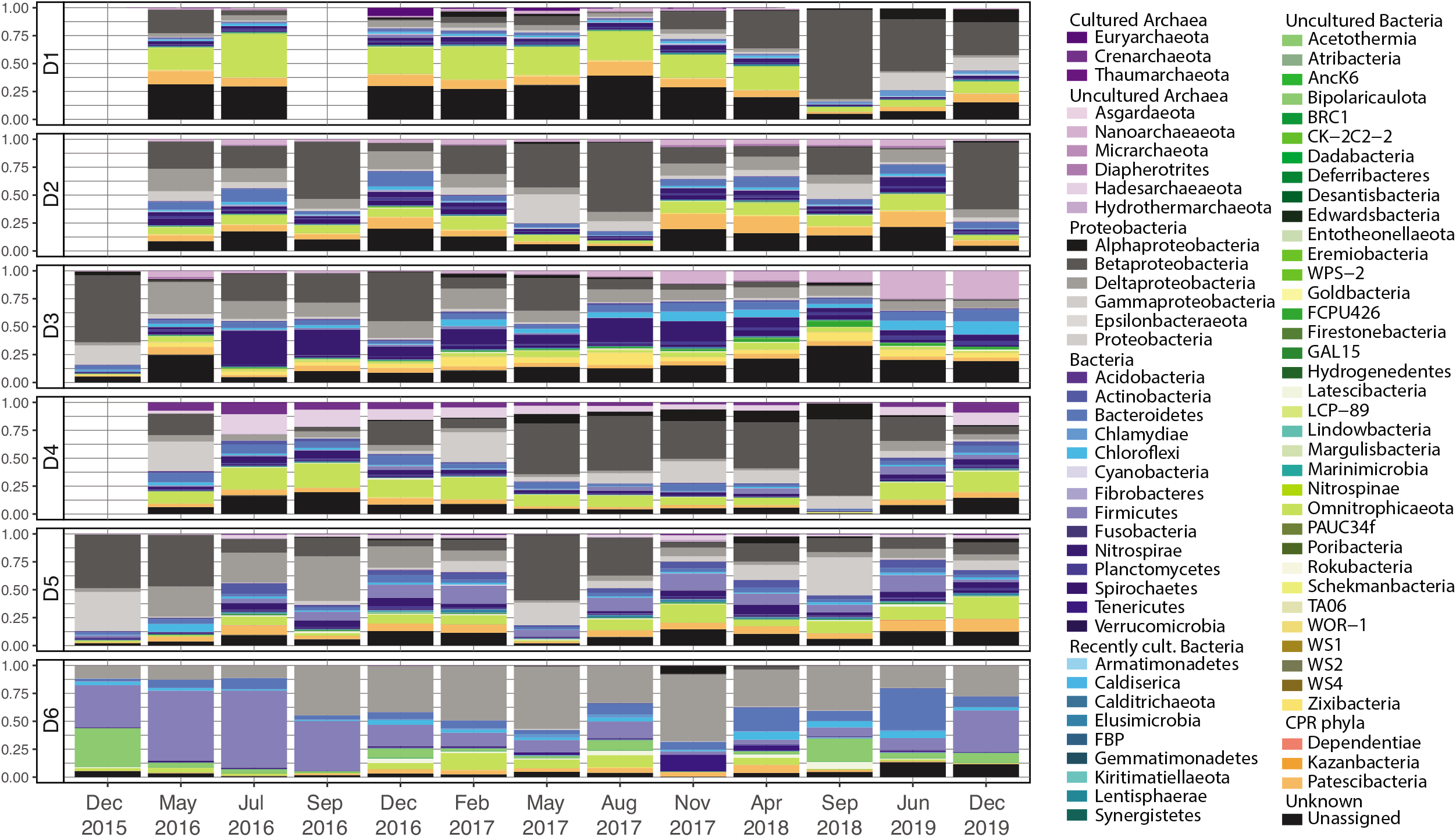

The most abundant OTUs (Sup. Fig. 1) at each site trend with phylum averages, but narrower taxonomic classifications are perhaps more meaningful. These OTUs are found within the Proteobacteria, Omnitrophicaeota, Firmicutes, Bacteroidetes (Ignavibacteriales, Lentimicrobiaceae), Chloroflexi, Nitrospirae, Patescibacteria (Pacebacteria), Zixibacteria, Acetothermia, Latescibacteria. Nanoarchaeaeota (Woesarchaeia), Bathyarchaeia and Hadesarchaeaeota as well as a number of discrete unassigned OTUs (#1, 25, 45). Proteobacteria include members of the Betaproteobacteria (*Sulfuricella, Sideroxydans*, Burkholderiaceae, Rhodocyclaceae, *Thiobacillus, Azospira* and *Ferribacterium*), Gammaproteobacteria (*Thiothrix*, Halothiobacillaceae), Deltaproteobacteria (Desulfobacteraceae, *Desulfovibrio, Smithella*, unclassified, Sva0485, *Desulfomicrobium, Desulfatiglans, Desulfobulbaceaea*). The Firmicutes include *Desulfurispora*, Thermoanaeobacteriaceae, Peptococcaceae, and an unclassified Clostridia. Nitrospirae distinctively includes Thermodesulfovibrionia and 4-29-1 classes. We discuss the ecological implications of these high abundance groups below.

Ecological diversity indexes (species richness, diversity, evenness) were used to assess differences across sites, both at single time points and for the 4-year average (Fig. 2, Sup. Table 1). D1 consistently showed the greatest species richness, followed by similar values for D2 and D3, then D4 and D5, and lastly D6. These patterns are similar for diversity metrics that incorporate abundance (Shannon and Simpson indexes) and phylogenic distance (Faith’s). D1 is consistently the most diverse and divergent; D2, D3, D4, and D5 are similar and intermediate, and D6 is consistently the least diverse and divergent. The control samples show high species richness and diversity with mean values comparable to D1 and a much larger range owing to the range of sample types (drilling fluid, mine ditch fluid from 800, 4100, and 4850 ft levels, as well as seeps and other borehole samples). All samples show very low evenness reflecting the presence of a few dominant taxonomic groups and diverse low abundance assemblages.

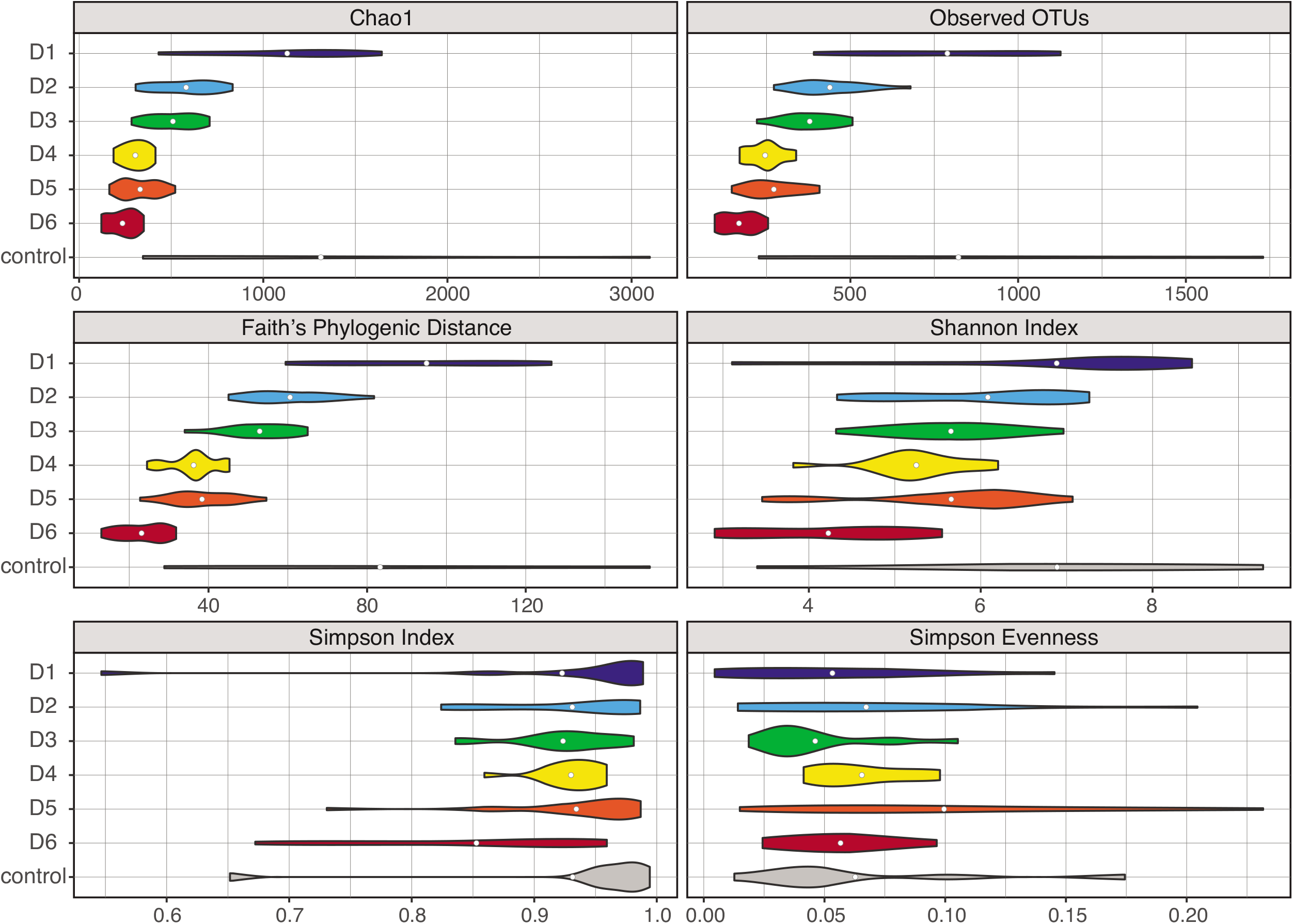

### 2.2 Beta diversity

The clear differences in taxonomy and diversity of DeMMO sites described above are quantified using the Bray Curtis dissimilarity coefficient, comparing taxonomic composition and relative abundances of each sample. These differences in community composition are plotted using nonmetric multidimensional scaling (NMDS) (Fig. 3), which shows clear grouping communities from each site over time. All time points from D6 plot closely together in the NW quadrant with 95% confidence ellipse including all but one point. Samples from D1 form a distinct cluster entirely encompassed by 95% confidence ellipses in the SW quadrant of the plot. Samples from D2 and D3 plot separately although proximally, in the NE quadrant with distinct and nonoverlapping 95% confidence ellipses. In contrast, samples from D4 and D5 plot together in the SW quadrant with overlapping 95% confidence ellipses containing approximately half of the points from each site. The same analysis, but including control samples (Sup. Fig. 2) illustrates strong separation between controls and borehole fluid samples, including two additional boreholes (DUSEL-B and LBNE-1) that were sampled irregularly over the course of the study. The control point plotting near D2 outliers consists of filtered natural seep fluids from the same level and may be geochemically similar to D2 fluids, thus hosting a similar community.

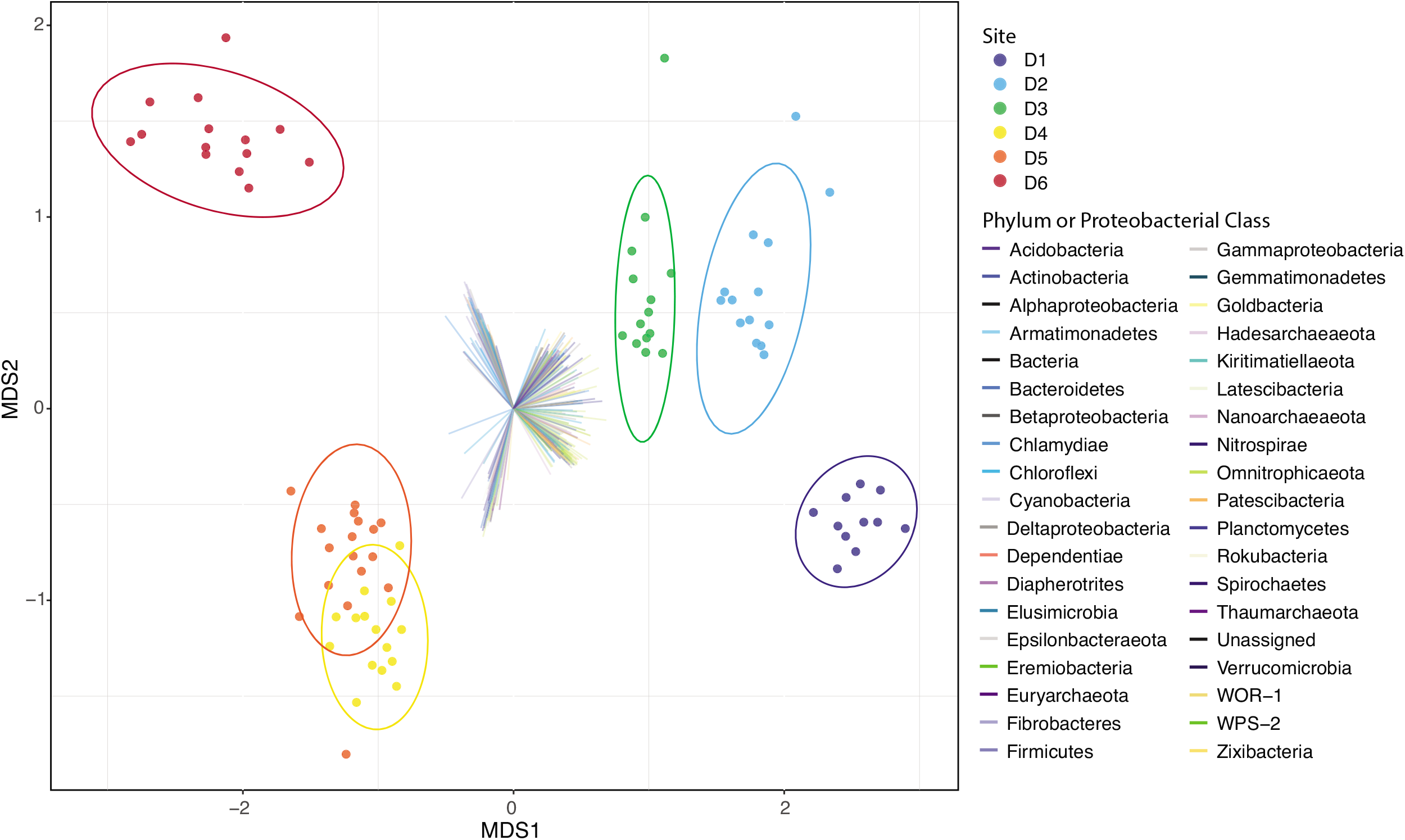

We can also ask which taxa correlate with different sites within this ordination framework. OTUs with significant (p <0.05) correlations are plotted on Fig. 3 using NMDS coordinates and colored by phylum. OTUs unique to a site ordinate in the direction of those samples, whereas those which are omnipresent have little statistical strength and are not plotted. For instance, the cluster of taxa pointing toward the D6 group includes a number of OTUs from the Firmicutes, Deltaproteobacteria, Chloroflexi, Tenericutes, and Bacteroidetes unique to that site. In contrast, the cluster ordinating toward the D1 group contains many OTUs from the Omnitrophicaeota, unassigned OTUs, and the Patescibacteria. While samples from D4 and D5 are largely overlapping, a cluster of OTUs ordinate strongly toward the D4 samples which includes Hadesarchaeaota, Crenarchaeota, Omnitrophicaeota, and Patescibacteria. Correspondingly, a relative few vectors ordinate toward the D5 cluster including Firmicutes, Bacteroidetes, and Chloroflexi. These differentiating taxa are distinguishing members of the communities on Fig. 1 and thus this approach confirms the statistical robustness of these observations. Control samples are differentiated from borehole samples by increased abundance of Proteobacterial groups, as well as Chloroflexi, Bacteroidetes, and Planctomycetes (Sup. Fig. 2).

### 2.3 Shared OTUs

One key question regarding the microbial ecology of the deep subsurface concerns the relative connectivity between sites, enabling exchange of fluids and microbial cells. To evaluate how similar the microbial populations are between DeMMO sites we evaluated how many OTUs were unique to a given site or shared between groups of two, three, four, five, or six sites (Fig. 4). In a system with limited hydrological connectivity, one would expect a high degree of unique OTUs at each site. In contrast, with greater connectivity, the same or similar OTUs could be shared across sites. With all other variables equal, one might expect a similar number of OTUs to be shared between each pair of sites (e.g. D1-D2), with fewer shared OTUs with each additional degree of grouping (e.g. D1-D2-D3) and geographic distance.

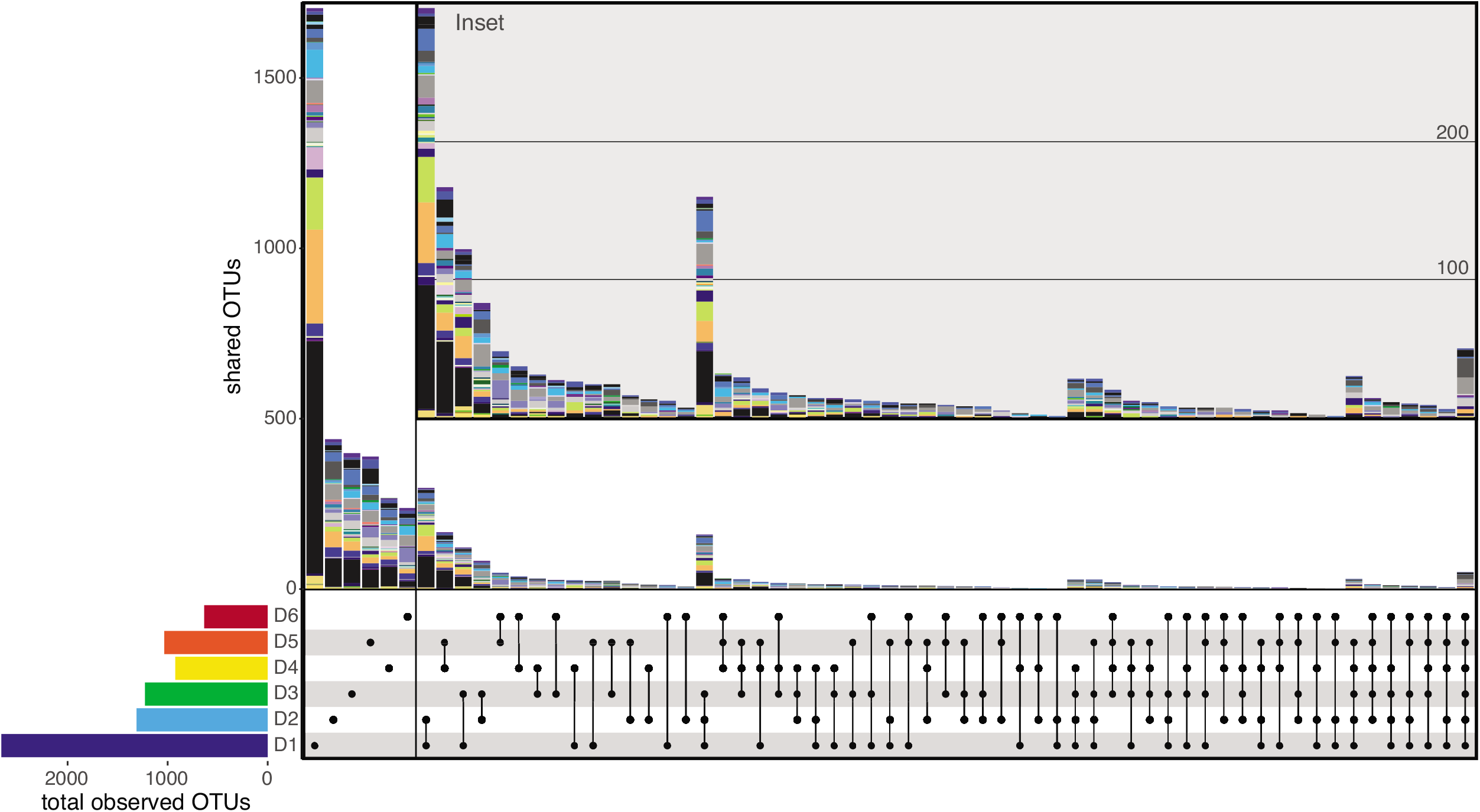

We observed a high percentage of unique OTUs in each site and a highly uneven distribution of shared OTUs between sites, consistent with limited hydrologic connectivity and microbial exchange. The number of OTUs unique to a site each parallels the total number of observed OTUs (bottom left color bars: D1>>D2>D3>D5>D4>D6), where the proportion of unique OTUs ranges from 64% at D1 to 29% at D4. The proportions for D2, D3, D5, and D6 are relatively similar (33% - 38%) suggesting that while the total diversity is variable, the level of site specificity remains consistent. D1 is exceptional, harboring the most observed OTUs (2664 total) and unique OTUs (64%), suggesting very high site-specific diversity. The stacked color bars illustrate the phylum-level classification of OTUs in each pairing and analysis of these unique OTUs is revealing. D1 harbors a vast number of unique unclassified OTUs as well as Patescibacteria and Omnitrophicaeota, and Nanoarchaeota, Deltaproteobacteria, and Chloroflexi at lower abundance. In contrast, the unique OTUs in the other five sites appear broadly similar at the phylum level, requiring that the divergence occurs at finer taxonomic resolution.

Comparing OTUs shared between sites can reveal degrees of potential connectivity and/or environmental similarity. Pairwise comparisons illustrate this point well with the highest number of shared OTUs shown by color bar height in Fig. 4 inset between D1 and D2 (297) then D4 and D5 (167), suggesting weighted similarity between shallow and deeper sites respectively. This is confirmed with triplets which are dominated by the D1-D2-D3 group (160) then the D4-D5-D6 (32). Clearly, depth is an important variable in determining the distribution of OTUs in the subsurface irrespective of temporal variation. Lastly, we must evaluate the 50 OTUs found across all sites. Optimistically, these taxa might represent subsurface specialists but, a more realistic interpretation would suggest that these are contaminants from DNA extraction and sequencing. Forty-four out of these 50 are also found in the control samples, suggesting contaminant origin. The remaining six are taxonomically related to uncultured strains within the Methanomassiliicoccales, Syntrophobacterales, Sva0485, Desulfobacteriales, Coriobacteriia, and Burkholderiaceae. The majority of these groups contain known anaerobes, sulfate reducers, and thermophiles, allowing for possible subsurface origin of these OTUs.

### 2.4 Microbial diversity in the context of site geochemistry

Parallel geochemical and microbial diversity data from our 4-year time series permits analysis of their potential interdependence. A correlation matrix between geochemical data, originally published in Osburn et al. (2019) and updated here, and microbial taxa distributions across DeMMO is illustrated in Fig. 5 and site specific correlograms are depicted in Sup. Fig. 4. While correlation does not imply causation, patterns apparent in these figures provide a starting point for ecological inquiry. In Fig. 5 clusters of taxa correlate with clusters of geochemical parameters, at the highest-level grouping geochemical characters into two large nodes, each broken up into two secondary nodes. Broadly, the leftmost nodes contain time variables, metals, water isotopes, SO_4_^2-^, DOC, dissolved H_2_ and CO_2_, ORP, Si and flow rate, where the rightmost nodes contain reduced gases, conductivity and associated ions (Cl, Li, Na), and pH in one sub-node and nutrients, temperature, and DIC and its isotopes in the other. Taxa distributions form four major nodes. One group diverges from all others forming two groups containing the Firmicutes and Alphaproteobacteria respectively. Next the Acetothermia group independently from the remaining taxa. The remaining taxa break out into two third-order nodes each with several subdivisions containing Deltaproteobacteria, Unassigned OTUs, and Omnitrophicaeota.

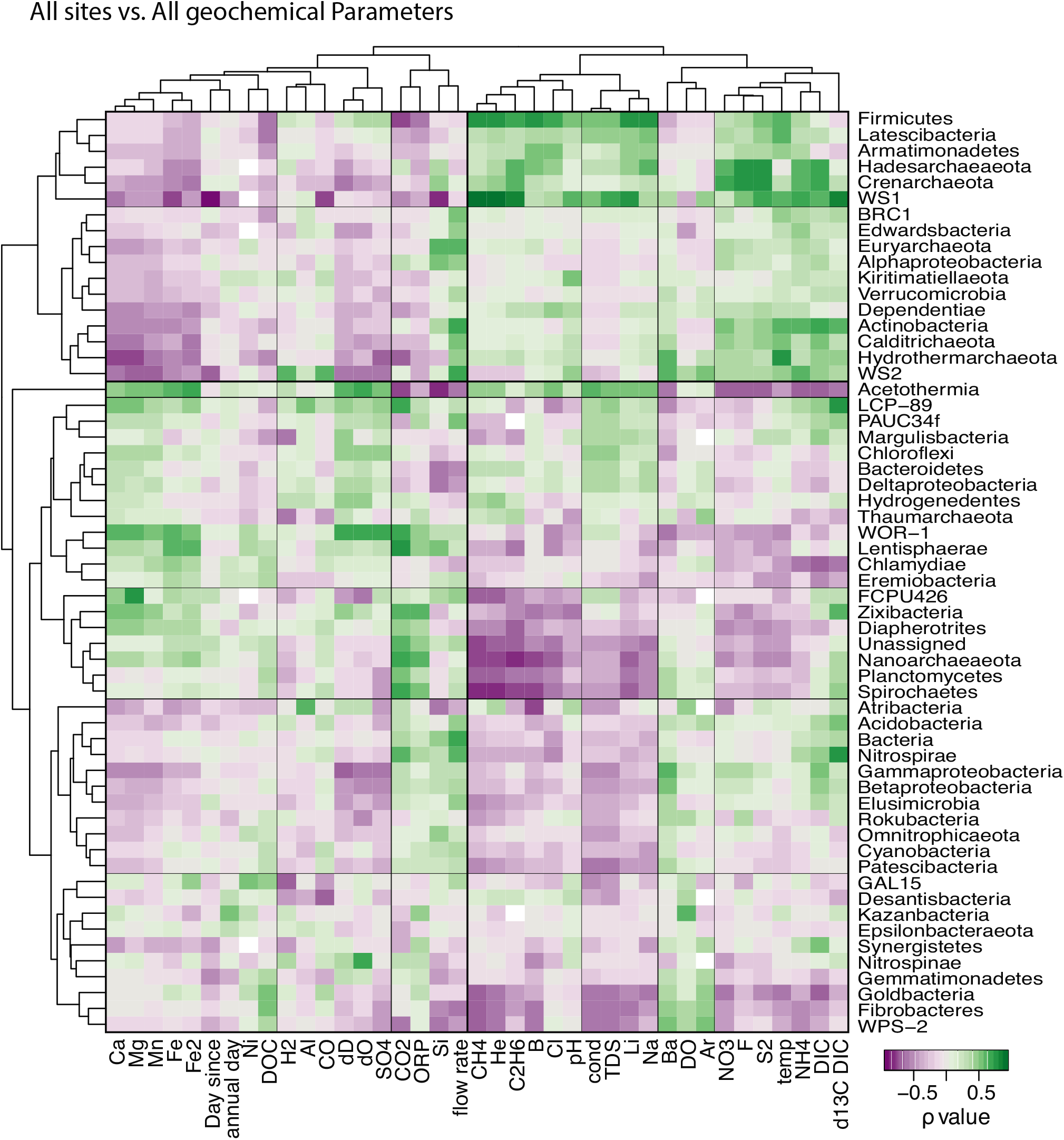

Comparing the geochemical and taxa groupings can inform environmental drivers. For instance, the group containing Firmicutes is strongly correlated with a geochemical cluster containing reduced gases (CH_4_, He, C_2_H_6_) as well as conductivity and associated ions, and another cluster containing nutrients (NO_3_^-^, NH_4_^+^, S^2-^, DIC) and temperature. These geochemical parameters characterize D5 and D6 and are where these taxa are enriched. Acetothermia correlates with the first cluster but not the second and is also associated with a metal cluster containing Ca, Mg, Mn, and Fe as well as SO_4_^2-^ and water isotopes, reflecting the geochemical composition of D6 alone. While, the grouping of high abundance taxa can be read off of Fig. 1, the distributions of less abundant taxa and their co-associations are observable here. For example, the Firmicutes cluster contains the Latescibacteria, Armatimonadetes, Hadesarchaeoeota, Crenarchaeota, and WS1, which again associate with anaerobic, high nutrient sites. Another strong cluster is between the Zixibacteria, Diapherotrites, Unassigned, Nanoarchaeaeota, Planctomycetes, and Spirochaetes. While little is known about the physiology of some of these taxa, this group is positively correlated with Ca, Mg, Mn, and Fe as well as CO2 and oxidation reduction potential (ORP). In aggregate, these associations suggest that these organisms are enriched in sites with high metal content, CO2 concentrations, and more oxidizing fluids. Since DeMMO sites themselves are geochemically and taxonomically distinct, microbial associations with particular sites likely drive the distributions in Fig. 5.

## 3. Discussion

### 3.1 Variability and stability in the dataset

The DeMMO network was designed to minimize bias during sampling, however, several significant environmental perturbations occurred over the course of this four-year experiment. First was establishment of the network in April-May of 2016, which altered passive fluid flow at each site. The degree of modification varied by site: D1 and D2 were fitted with standard drain packers; D3, D4, and D5 were reamed out and custom packers were installed; and D6 was left unmodified (see Osburn *et al*. (2019) for details). Another perturbation was the installation of flow-through colonization reactors beginning in July 2016 (Casar *et al*., 2020). Because reactors were fitted on side ports at D1-D5 while still allowing passive fluid outflow from the sites, the installations resulted in only minor geochemical changes but at D6, the installation initiated open flow from a previously closed system. This change affected the fluid geochemistry and ultimately drained the borehole by May 2017. After the site was returned to a closed system and refilled, the measured geochemical parameters returned to pre-perturbation values. This supports our hypothesis that these boreholes are fed by slowly migrating subsurface fluids which are only minimally affected by mine activity.

Based on the site histories, differences in flow patterns, geochemical parameters, and microbial diversity, the microbial communities of DeMMO sites may exhibit different degrees of variability over time which can be assessed in different ways (Hughes *et al*., 2001; Martínez-Martínez *et al*., 2006). For example, population variance describes the spread of a given population around its mean from a number of sampling points. We calculated this value for each taxon, but given the large differences in population means, variance is primarily dependent on the abundance of a given taxon. To minimize this effect and to emphasize changes within the less abundant taxa, we normalized the variance by the average taxon abundance, producing a variance to abundance ratio (var:abu). This measure is conceptually similar to dispersion, but uses variance instead of standard deviation. Under this framework, a group with a population variance equal to its mean abundance would have a var:abu ratio of 1. Groups falling significantly above this threshold (>1.5) are considered highly variable, whereas those falling significantly below (<0.5) are stable through time. We will use these thresholds (>1.5, 1.5 to 0.5, and <0.5) to demarcate high, medium, and low taxa variability.

Table 1 catalogs the number of OTUs and Phyla (and Proteobacterial Classes) that are included in the aforementioned variability groups. We see that the number of highly variable phyla is small compared to low variability phyla (e.g. D2, >1.5 = 5, <0.5 = 32). When viewed at the OTU level, this trend is even more dramatic with only 12 to 32 OTUs falling in the high variability groups compared with hundreds to thousands in the low variability groups (depending on the site). Thus, the vast majority of taxa (85 to 99% of OTUs) within a site maintained stable population abundance over this four-year experiment. That said, the relative abundances of these groups vary markedly between sites. For instance, in D1 the highly variable taxa make up only 32% of average taxa abundance, leaving the majority of the community in at medium (9%) to low (59%) variability. In contrast, at D6 highly variable OTUs make up 77% of the average OTU abundance with only 15% of OTUs in the stable category (Table 1).

**Table 1:** The relative variability and abundance of OTUs in each site.

|  | **Obs. OTUs** | **# of Phyla** |  |  | **# of OTUs** |  |  | **% ave. abundance** |  |
| --- | --- | --- | --- | --- | --- | --- | --- | --- | --- |
|  |  | **>1.5** | **1.5-0.5** | **<0.5** | **>1.5** | **1.5-0.5** | **<0.5** | **>1.5** | **<0.5** |
| **D1** | 2664 | 7 | 3 | 45 | 12 | 19 | 2633 | 32.3 | 58.7 |
| **D2** | 1311 | 5 | 11 | 32 | 13 | 29 | 1269 | 38 | 43.3 |
| **D3** | 1227 | 7 | 9 | 36 | 23 | 31 | 1173 | 50.6 | 33 |
| **D4** | 924 | 8 | 6 | 30 | 29 | 30 | 865 | 63.6 | 23.4 |
| **D5** | 1034 | 11 | 5 | 37 | 32 | 46 | 956 | 51.7 | 27.3 |
| **D6** | 635 | 11 | 2 | 25 | 24 | 28 | 539 | 76.8 | 15.2 |

#### 3.1.1 Variable Taxa

In order to delve deeper into these highly variable phyla, we have plotted their abundance through time from each site (Sup. Fig. 3 and Fig. 1). While all sites show large changes, they express different styles and degrees of variability with some corresponding to known environmental triggers and others appearing more random. For instance, at D1 a broad decrease in Omnitrophicaeota abundance is paired with a dramatic increase in Betaproteobacteria starting in Nov. 2017, temporally consistent with the installation of flow through experiments at this site (Casar et al., 2020). A similar mechanism is likely at play at D4 where Omnitrophicaeota decrease and Betaproteobacteria increase starting in May 2017. At D3 a broad decrease in Proteobacteria and Nitrospirae is paired with an increase in Chloroflexi and Nanoarchaeaeota, although this corresponds with no known perturbation. At D2 the variability appears scattered rather than as a long-term trend with periodic spikes in Beta- and Gammaproteobacteria as well as Bacteroidetes and Patescibacteria. Similarly, at D5 spikes in Beta-, Delta-, and Gammaproteobacteria, Firmicutes, Chloroflexi, and Omnitrophicaeota occur throughout. D6 exhibits the most striking changes with time. Initially Acetothermia and Firmicutes are at near equal abundance before Acetothermia drop and Firmicutes rise, only to be replaced by the Deltaproteobacteria. Patescibacteria follow Deltaproteobacteria at lower abundance. Subsequently Bacteroidetes rise followed by revival of Acetothermia and Firmicutes populations. These changes have concrete physical triggers as this manifold was opened in July 2016, drained by December 2016, and subsequently returned to a closed system. These physical changes appear to have had a dramatic effect on the abundance of variable taxa.

Paired microbial and geochemical datasets are integral for identification of environmental triggers for microbial change. To this end, we repeat the correlation analysis shown in Fig. 5 for samples from each site (Sup. Fig. 4). Here, taxa and geochemical parameters are grouped by parameters that vary together within a given site, thereby revealing covariance. Temporal trends can be seen by comparison of experimental duration (‘Day since’ column) with microbial taxa for each panel in Sup. Fig. 4. These correlations are consistent with those displayed in Sup. Fig 3 but reveal the behavior of minor taxa and paired geochemical changes. For instance, at D1 we see a negative correlation with time (decreases) for Omnitrophicaeota, Gemmatimonadetes, and Actinobacteria (among others) and a positive correlation (increases) with Proteobacterial classes. Time correlates with Si, B, Mn, Fe, conductivity, sulfide and DIC, which could be physiologically related to taxa abundances. At D3 we see increases in Chloroflexi, Nanoarchaeaeota, Planctomycetes, Unassigned, and FCPU426 and decreases in Goldbacteria, Gammaproteobacteria, Betaproteobacteria, Cyanobacteria, and Nitrospirae, corelating to changes in the concentration of SO_4_^2-^, NO_3^-^_, metals, and conductivity.

Microbial variability lacking a strong temporal trend will correlate instead with geochemical parameters (Fig. Sup. 4). Here we discuss the taxa that corelate with the most variable geochemical parameters in each site as identified in Osburn et al. (2019). Geochemically, D1 shows minor changes in Fe^2+^, CO_2_, and DOC concentrations and higher Fe^2+^ corresponds to higher Omnitrophicaeota and Epsilonproteobacteria and lower Betaproteobacteria and Armatimonadetes abundances, suggesting a potential relationship with iron. At D2, NO3^-^ varies most, correlating to increases in Alphaproteobacteria and decreases in Spirochaetes. D3 is temporally more variable particularly in ORP, δ^13^C_DIC_, NH_4_^+^, and Fe^2+^ and correspond to increases in Beta- and Gammaproteobacteria and Goldbacteria and decreases in Chloroflexi, Planctomycetes, FCPU426, and Bacteroidetes. D4 and D5 both have high and fluctuating sulfide concentrations. At D4 higher sulfide corresponds to higher Armatimonadetes, BRC1, and Actinobacteria and lower Zixibacteria, Firmicutes, and Latescibacteria abundances. In contrast, at D5 higher sulfide corresponds to higher WOR-1and Nanoarchaeaeota, and lower Armatimonadetes abundances. Large geochemical variations occur at D6 including changes in Fe^2+^ concentration and ORP which correspond to increases in Delta- and Betaproteobacteria, Patescibacteria, and Planctomycetes and decreases in Firmicutes, Nitrospirae, Acetothermia, and Bacteroidetes. Reduced gases (He, CH_4_, and C_2_H_6_) cluster together and correlate to increases in Nitrospirae and Bacteroidetes whereas increases in DOC correspond to increases in Bacteroidetes and decreases in BRC1. While these correlations should not be interpreted as causative, many of the taxa described here are uncultured. While a physiological interpretation is not possible here, this analysis identifies patterns which can subsequently be tested with genome analysis or targeted culture-based enrichments.

#### 3.1.2 Stable Taxa

While variable taxa reveal subsurface population dynamics, perhaps the most critical observation of this this study is stability: the vast majority of OTUs (85% and 99%) are essentially invariant over this four-year experiment. While these stable OTUs are often at low abundance, their ubiquitous presence warrants further inquiry and may have implications for subsurface biospheres elsewhere. Comparison of OTU proportion and relative abundance of variable and stable groups emphasizes the sheer number of stable taxa and reveals strong differences in their taxonomy (Fig. 6). Stable OTUs are found widely across the tree of life and are evenly distributed across sites in both proportion of observed OTUs (Fig. 6A) and relative abundance (Fig. 6B). In particular they contain containing many Nanoarchaeaeota, Proteobacteria, Acidobacteria, Actinobacteria, Chloroflexi, Firmicutes, Nitrospirae, Planctomycetes, Omnitrophicaeota, Patescibacteria, and Unassigned groups. In contrast, variable OTUs occur in a limited number of phyla (Proteobacteria, Nitrospirae, Acetothermia, Bacteroidetes, Hadesarchaeaeota, Nanoarchaeaeota, and Crenarchaeota) and in low numbers with uneven abundance between sites, often appearing at high abundance in only one site. Omnitrophicaeota and Firmicutes are found in high numbers and abundance in both stable and variable groups with uneven site distributions.

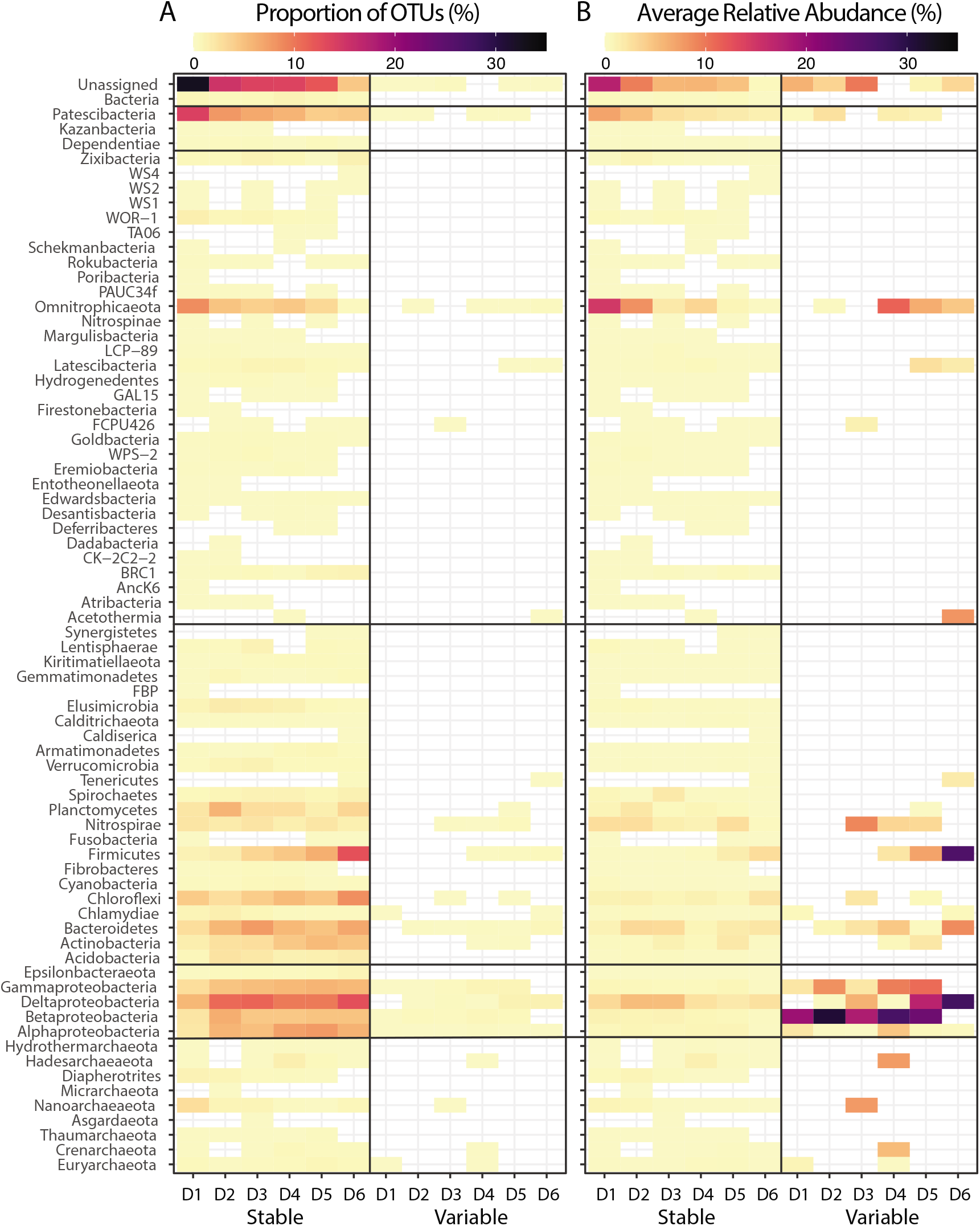

The site distribution of stable and variable OTUs correlates with geochemical variability. D6, the most dynamic site, has the lowest proportion of stable OTUs while D1, the most stable, has the highest. Proteobacterial classes are evenly distributed across the stable OTUs, but are far less abundant than their variable cousins, suggesting the variable metabolic potential of Proteobacterial groups may be ideal for responding to environmental perturbation. In contrast, the diverse membership of the stable OTUs, particularly within the candidate phyla, may suggest strong niche partitioning and metabolic fragmentation in this core subsurface microbiome.

### 3.2 Ecological significance to the subsurface biosphere

The subsurface biosphere has unique physical and geochemical conditions that influence the taxa capable of inhabiting this realm. While there is no consensus in the literature thus far on which taxa constitute a typical subsurface biome, locations worldwide suggest the presence of methanogens, sulfate reducers, spore formers, and vast diversity of novel clades including CPR taxa (Chivian *et al*., 2008; Castelle *et al*., 2013; Hug *et al*., 2015; Purkamo *et al*., 2016; Anantharaman *et al*., 2016; Lau *et al*., 2016; Momper, Kiel Reese, *et al*., 2017; Kadnikov *et al*., 2017; Probst *et al*., 2018; Purkamo *et al*., 2020). Indeed, we see many microbial taxa found previously in subsurface environments including Proteobacteria, Firmicutes, Chloroflexi, Nitrospirae, Archaea, and numerous uncultivated lineages inhabiting DeMMO sites. Interrogation of the membership of these groups can provide a glimpse into the potential metabolisms and adaptations that contribute to their subsurface fitness.

Proteobacteria are diverse and abundant in DeMMO and dominate the variable OTUs. Metabolic plasticity and diversity of many Proteobacteria may contribute to this distribution (Garrity and Kuever). Proteobacterial classes vary with depth, with Betaproteobacteria in shallow sites, Beta-, Delta-, and Gammaproteobacteria in mid-depth, and mostly Deltaproteobacteria at depth. Many abundant OTUs come from well-classified genera with distinctive metabolic capability. For instance, the Betaproteobacteria *Sideroxydans*, *Sulfuricella*, and *Thiobacillus* are prevalent in D1, D2, and D3, whereas D4 and D5 contain members of the *Azospira*, *Ferribacterium*, unclassified Burkholderiaceae, and *Thiobacillus*. Prevalent Gammaproteobacteria include *Thiothrix* and Halothiobacillaceae. Deltaproteobacteria across all sites belong to sulfate-reducing groups including *Desulfovibrio*, *Desulfomicrobium*, and unclassified Desulfobulbaceaea. In total, this distribution suggests that metal-cycling and oxidative sulfur metabolisms dominate in shallow sites and sulfate reduction takes over at depth.

Firmicutes are often suggested to be subsurface specialists due to their spore-forming capabilities and common anaerobic metabolisms. The most famous is *Desulforudis audaxviator* which was found in South African gold mines and for which genome sequencing revealed diverse metabolic capabilities fueled primarily by autotrophic sulfate reduction (Chivian *et al*., 2008; Lau *et al*., 2016). Firmicutes are abundant in D4, D5, and D6 (increasing in that order). The most abundant OTUs are unclassified Peptococcaceae (the same family as *Desulforudis*), but also *Desulfurispor*a, Thermoanaerobacteriaceae, and unclassified Clostridia. While the characterized *Desulfospora* have similar metabolic capabilities to *Desulforudis* (Kaksonen *et al*., 2007) suggesting sulfate-reduction at DeMMO, the remaining Firmicutes are too divergent to speculate on their characteristics beyond likely being anaerobic spore-formers.

Numerous Chloroflexi are found in the stable OTUs, consisting almost entirely of uncultivated Anaerolineales and *Dehalococcoida*. *Dehalococcoida* has been widely observed in the marine subsurface (Wasmund *et al*., 2014) and in aquifer sediments (Hug *et al*., 2013). Subsurface Anaerolineales have been found in hydrothermal sediments off of Japan where they are suggested to be mediating anaerobic heterotrophy or mixotrophy (Fullerton and Moyer, 2016) as well as in a Siberian deep aquifer where they are implicated in aerobic and anaerobic respiration and fermentation (Kadnikov *et al*., 2018). The Anaerolineales contain complete Acetyl-CoA carbon fixation pathways, suggesting the potential for *in situ* carbon fixation. Similar roles are possible for the Chloroflexi at DeMMO, but must be confirmed with additional approaches.

Nitrospirae are common subsurface residents and ubiquitous members of all DeMMO sites, pervading the stable OTUs and containing abundant variable members in D3, D4, and D5. The majority of these Nitrospirae are uncultured orders within the Thermodesulfovibrionia. This class has one lineage with cultivars (*Thermodesulfovibrio*) comprised of sulfur cycling thermophiles isolated from the terrestrial subsurface (Frank *et al*., 2016). However, uncultivated DeMMO strains were found to colonize coupons of manganese oxide at low temperature, suggesting their potential role in mineral metabolism and biofilm formation (Casar *et al*., 2020).

Surprisingly, Chlamydiae are ubiquitous across DeMMO. While these bacteria are known as obligate intracellular eukaryotic parasites, increasing evidence suggests that they infect free-living amoebae within environmental waters (Corsaro *et al*., 2009) and have been identified in shallow sedimentary aquifers (Hug *et al*., 2015). We find a diversity of Chlamydiae classes in both stable and variable populations. If these are eukaryotic parasites, this would imply widespread colonization of deep subsurface groundwaters by amoebae, consistent with observations of other eukaryotes (Borgonie *et al*., 2011; 2015).

Archaea feature strongly in the variable and stable populations including both cultivated and uncultivated lineages. Euryarchaeota are found at low levels ubiquitously including members of Methanomicrobia and Thermoplasmata. Where these OTUs are sufficiently classified, most derive from putative methanogenic groups. Crenarchaeotal OTUs are members of the Bathyarchaeia which have been found widely in marine sediments (Feng *et al*., 2019). Methanogenic archaeal groups are commonly observed in both the marine and continental subsurface. Among the uncultured archaea, Nanoarchaeota, Diapherotrites and Hadesarcharchaeota are consistently present and abundant in certain sites. Nanoarchaeaeota are prevalent in D3 variable OTUs whereas Hadesarchaeaota are abundant D4. The Nanoarachaeaota are members of DPANN and share a hypothesized reduced metabolic capacity with members of the CPR (Castelle and Banfield, 2018). Hadesarchaeaota, were originally described from a South African gold mine and are genomically implicated to perform CO oxidation, carbon fixation, heterotrophy, and nitrogen-based metabolisms (Baker *et al*., 2015).

Metabolic and ecological evaluation of uncultivated lineages is very challenging and requires analyses beyond the scope of this work. However, Candidate Phyla are very common within DeMMO, suggesting their ecological importance. In addition to the archaea described above, we commonly observe abundant Omnitrophicaeota and Patescibacteria, as well as numerous low abundance lineages including Acetothermia, Latescibacteria, WOR-1, BRC1, and Zixibacteria. Omnitrophicaeota are particularly abundant in the stable OTUs at D1, D2, and D4 and variable OTUs of D4, D5, and D6 (Fig. 6). This group (originally Omnitrophica) was defined in Rinke et al. (2013) and suggested to perform carbon fixation via the Acetyl-CoA pathway, a hypothesis that was confirmed in Momper et al. (2017), who also found evidence for methane oxidation, as well as nitrate and sulfate reduction (Momper, Jungbluth, Lee, and Amend, 2017b). CPR taxonomy is rapidly evolving and contradictory with most references referring to the CPR superphylum with many phyla (35, (Brown *et al*., 2015)) at odds with the Patescibacteria phylum used in the Silva classification applied here (Hug *et al*., 2016; Castelle and Banfield, 2018). Regardless, these organisms are common subsurface residents identified widely in shallow groundwaters, united by a tendency toward extremely reduced genomes and streamlined metabolic capabilities (Brown *et al*., 2015; Anantharaman *et al*., 2016; Probst *et al*., 2017; Castelle and Banfield, 2018; Tian *et al*., 2020). We decline to speculate on the physiology of the other uncultivated phyla and direct the reader to our forthcoming metagenomic studies.

## 4. Conclusions

Here we present a unique record of paired subsurface microbial diversity and geochemistry extending 1.5 km through the crust, collected for over four years. This record is the first of its kind and illuminates key aspects of subsurface diversity and temporal dynamics. We see decreasing alpha diversity with depth and distinct beta diversity between sites strongly driven by distinct geochemistry. This dataset reveals variable and stable populations, uniquely describing dynamics within microbial groups within a backdrop of a stable core microbiome. The generally abundant variable OTUs are composed of Proteobacteria, Omnitrophicaeota, Patescibacteria, Zixibacteria, Acetothermia, Nitrospirae, and Firmicutes. Whereas the stable core comprises 88-99% of observed OTUs and covers a rich swath of cultured and uncultured microbial taxa. OTU abundance relates to geochemical stability of each site, with the most geochemically stable sites harboring the most stable populations.

This pattern of contrasting dynamics and stability may characterize the terrestrial subsurface biosphere where populations must persist over long time periods but also be poised to exploit periodic resource opportunities. Together, these taxa contribute to a diverse subsurface ecosystem with much in common with previously characterized locations, suggesting an emerging group of core subsurface specialists. While no two subsurface sites are the same, there are common threads including an enrichment of Proteobacteria, Firmicutes, unclassified groups, uncultured bacterial and archaeal phyla, and members of the CPR. Metabolically these groups have the potential for widespread carbon utilization and fixation, sulfur and metal-based metabolisms, and methanogenesis.

## 5. Experimental Procedures

Samples were collected from the DeMMO network between on fourteen trips between Dec. 2015 and Dec. 2019. Owing to facility logistics, each sampling campaign occurred over 4 days. Geochemical samples were collected as described in Osburn et al. (2019). In short, filtered fluid was collected for IC, ICP, DIC, DOC analysis, whereas redox sensitive ions were measured on site using field spectrophotometers. Samples of borehole manifold effluent for DNA sequencing were taken using Sterivex brand 0.22 μm filters while monitoring filtrate volume (between 0.5 and 1 L). The filters were placed in a sterile falcon tubes on dry ice prior to transport back to the lab and storage at −80°C. For samples taken before borehole modification and some control samples, fluid was collected in a sterile 60 mL syringes and manually filtered.

DNA was extracted using either the MoBio Sterivex kit or MoBio PowerSoil DNA Isolation kit following the manufacturer instructions. Whole genomic DNA was sent to the Environmental Sample Preparation and Sequencing Facility at Argonne National Laboratory for 16S rRNA gene amplicon sequencing of the V4 hypervariable region using 516F/806R universal primers on an Illumina MiSeq platform following manufacturer and facility protocols.

Paired-end reads were joined using PEAR v 0.9.8 (J. Zhang *et al*., 2014) and demultiplexed with QIIME v 1.9.1 (Caporaso, 2010). Sequences were dereplicated and binned into operational taxonomic units OTUs at a threshold of 97% similarity and chimeric sequences were removed using USEARCH v 11.0.667 (Edgar, 2010). While we acknowledge that ASVs are risign as the preferred sequence analysis technique, the long term, multi-batch nature of these data and higher taxonomic resolution requires and permits the use of OTUs. Reads were rarefied to a depth of 6,500. OTUs were assigned taxonomy using the SILVA132 database (Quast *et al*., 2012). “Unassigned” OTUs were compared to DeMMO metagenome-assembled genomes (MAGs) for further assignment following methods described in Casar et al. (2020). We performed statistical analyses on the rarefied OTU table using QIIME and the Vegan v 2.5-5 (Oksanen *et al*., 2019) and Ecodist v 2,9,1 (Goslee and Urban, 2007) packages in R. Alpha diversity was calculated using QIIME. Non-metric multidimensional scaling (NMDS) ordination was carried out using Vegan on communities at the OTU level using the metaMDS function Bray-Curtis metric with default parameters and a dimension size of 2.

## Acknowledgements

Working in the deep subsurface requires the cooperation of many institutions and individuals. We would like to thank SURF staff including Jaret Heiss, Tom Regan, Kathy Hart. Sequencing analyses were assisted by Sarah Owens and Stephanie Greenwald. Funding for this work was provided by the NASA Astrobiology Institute Life Underground (PI JPA), a NASA Exobiology grant to MRO, a Packard foundation fellowship to MRO, and a CIFAR fellowship to MRO, and a NASA NESSF to MRO and CPC. All code and corresponding data used in this study are available at github.com/CaitlinCasar. The authors have no conflict of interests to declare. Sequence data are available in the NCBI GenBank database under accession KDHR00000000.

