## Supplemental for "Contrasting Variable and Stable Subsurface Microbial Populations: an ecological time series analysis from the Deep Mine Microbial Observatory, South Dakota, USA"

**Supplementary Information**

**Tables:**

**Supplementary Table 1:** Alpha Diversity Metrics

|  | **Chao 1** | | | **Observed OTUs** | | |
| --- | --- | --- | --- | --- | --- | --- |
|  | min | max | mean | min | max | mean |
| **D1** | 432 | 1644 | 1130 | 391 | 1127 | 789 |
| **D2** | 307 | 834 | 581 | 272 | 679 | 439 |
| **D3** | 285 | 709 | 509 | 221 | 506 | 379 |
| **D4** | 187 | 415 | 305 | 170 | 338 | 246 |
| **D5** | 164 | 522 | 331 | 146 | 408 | 272 |
| **D6** | 121 | 351 | 235 | 96 | 255 | 168 |
| **controls** | 347 | 3099 | 1313 | 227 | 1730 | 822 |
|  | **Phylogenetic Distance** | | | **Shannon Index** | | |
|  | min | max | mean | min | max | mean |
| **D1** | 59 | 127 | 95 | 3.1 | 8.5 | 6.9 |
| **D2** | 45 | 82 | 61 | 4.3 | 7.3 | 6.1 |
| **D3** | 34 | 65 | 53 | 4.3 | 7.0 | 5.7 |
| **D4** | 24 | 45 | 36 | 3.8 | 6.2 | 5.2 |
| **D5** | 23 | 55 | 38 | 3.5 | 7.1 | 5.7 |
| **D6** | 13 | 32 | 23 | 2.9 | 5.5 | 4.2 |
| **controls** | 29 | 151 | 83 | 3.4 | 9.3 | 6.9 |
|  | **Simpson Index** | | | **Simpson Evenness** | | |
|  | min | max | mean | min | max | mean |
| **D1** | 0.55 | 0.99 | 0.92 | 0.005 | 0.145 | 0.053 |
| **D2** | 0.82 | 0.99 | 0.93 | 0.014 | 0.204 | 0.067 |
| **D3** | 0.84 | 0.98 | 0.92 | 0.019 | 0.105 | 0.046 |
| **D4** | 0.86 | 0.96 | 0.93 | 0.041 | 0.098 | 0.065 |
| **D5** | 0.73 | 0.99 | 0.93 | 0.015 | 0.232 | 0.100 |
| **D6** | 0.67 | 0.96 | 0.85 | 0.024 | 0.096 | 0.057 |
| **controls** | 0.65 | 0.99 | 0.93 | 0.013 | 0.174 | 0.063 |

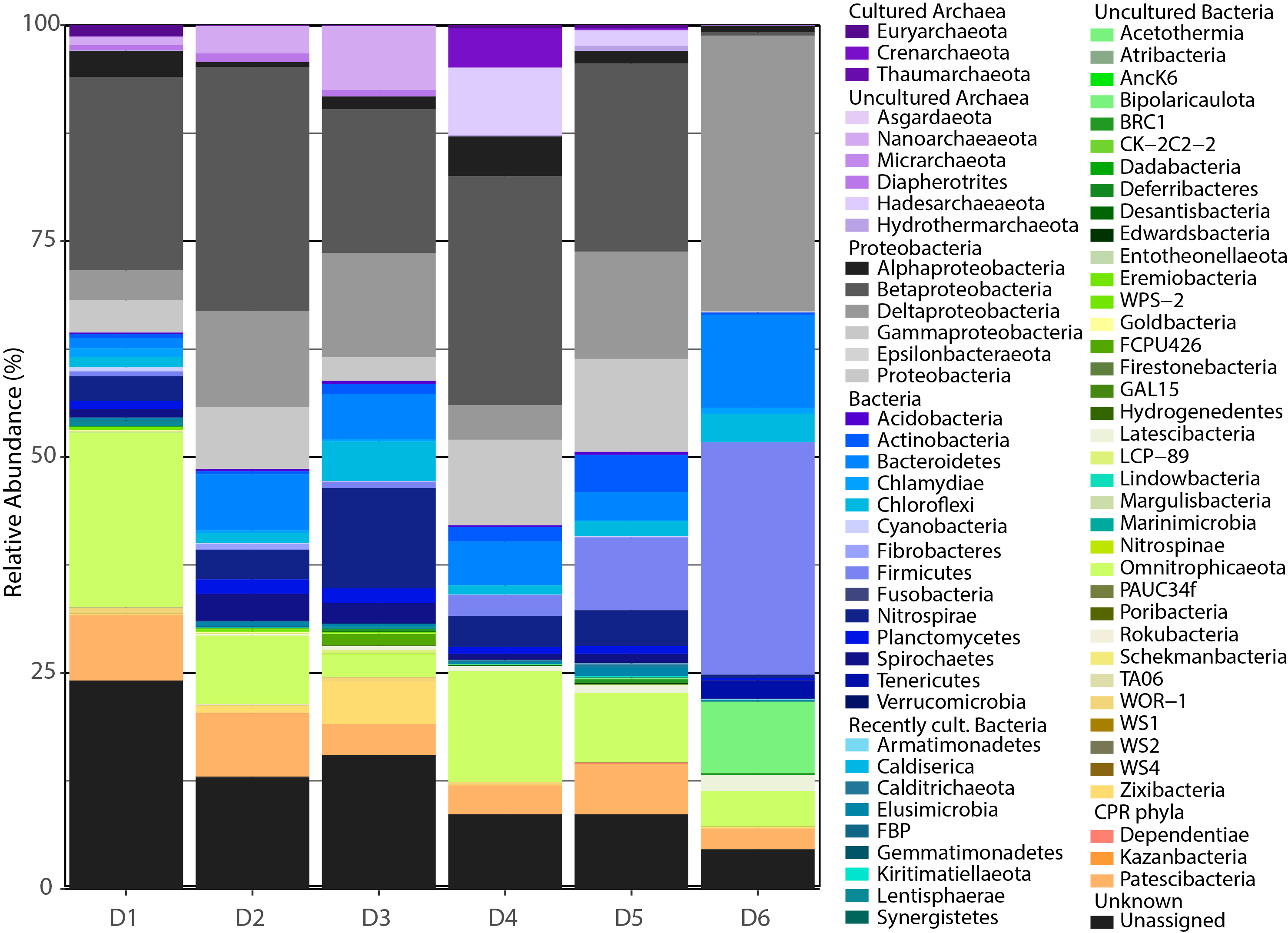

**Supplementary Figure 1**: Phylum-level abundance of microbial taxa in each site on average. Proteobacterial classes are divided in grey scale. Phylogeny was assigned with reference to the Silva 132 database as well as an internal GTDBtk-assigned genome database.

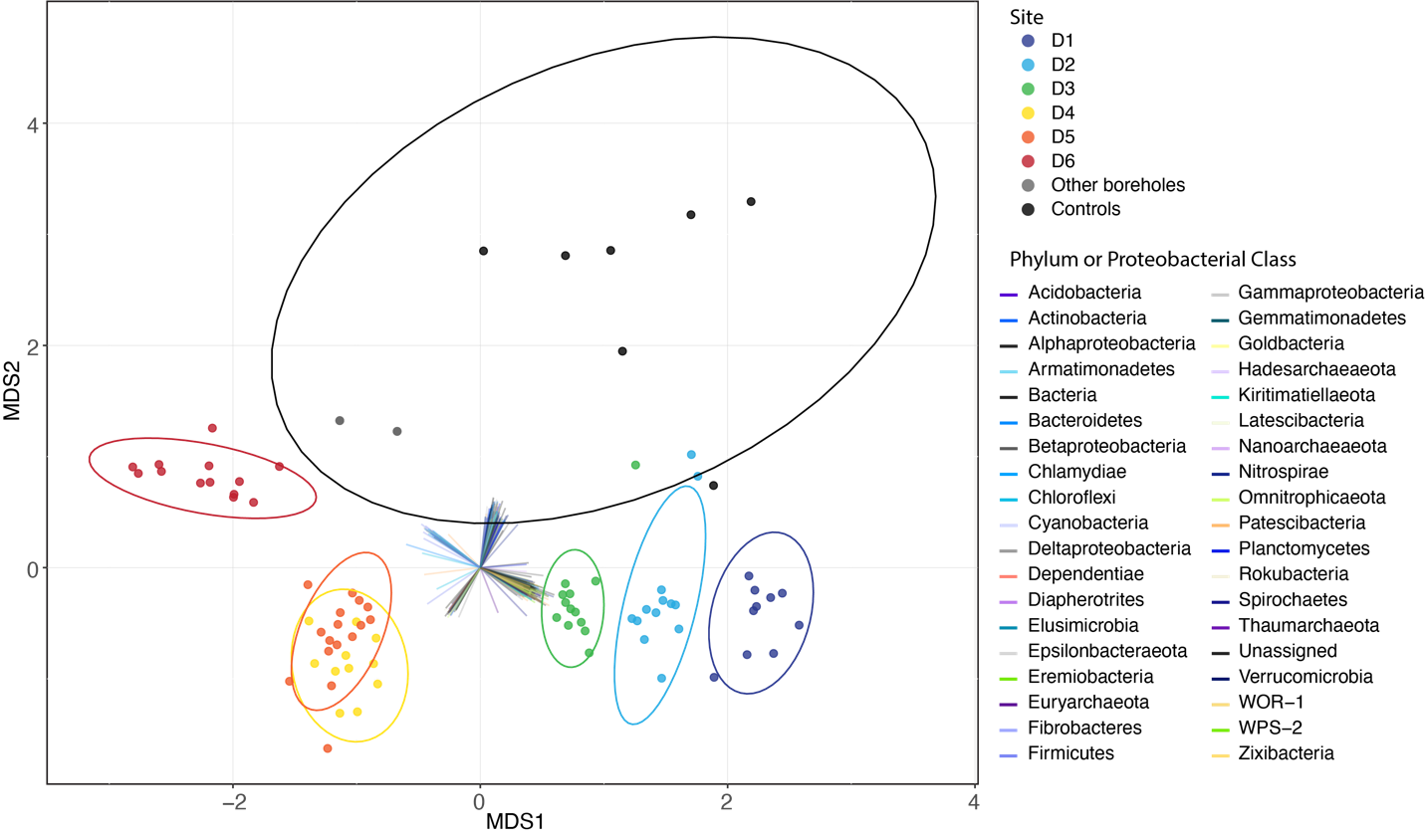

**Supplementary Figure 2:** NMDS ordination of all time points from each DeMMO site and control samples based on Bray Curtis Dissimilarity. 95% confidence ellipses are shown for each site and show clear separation between all but the D4-D5 pair. OTUs with statistically significant correlations (p <0.05) are plotted on the same ordination, illustrating the factors responsible for separation of sample groups. Please see the supplementary HTML version to explore specific OTU distributions.

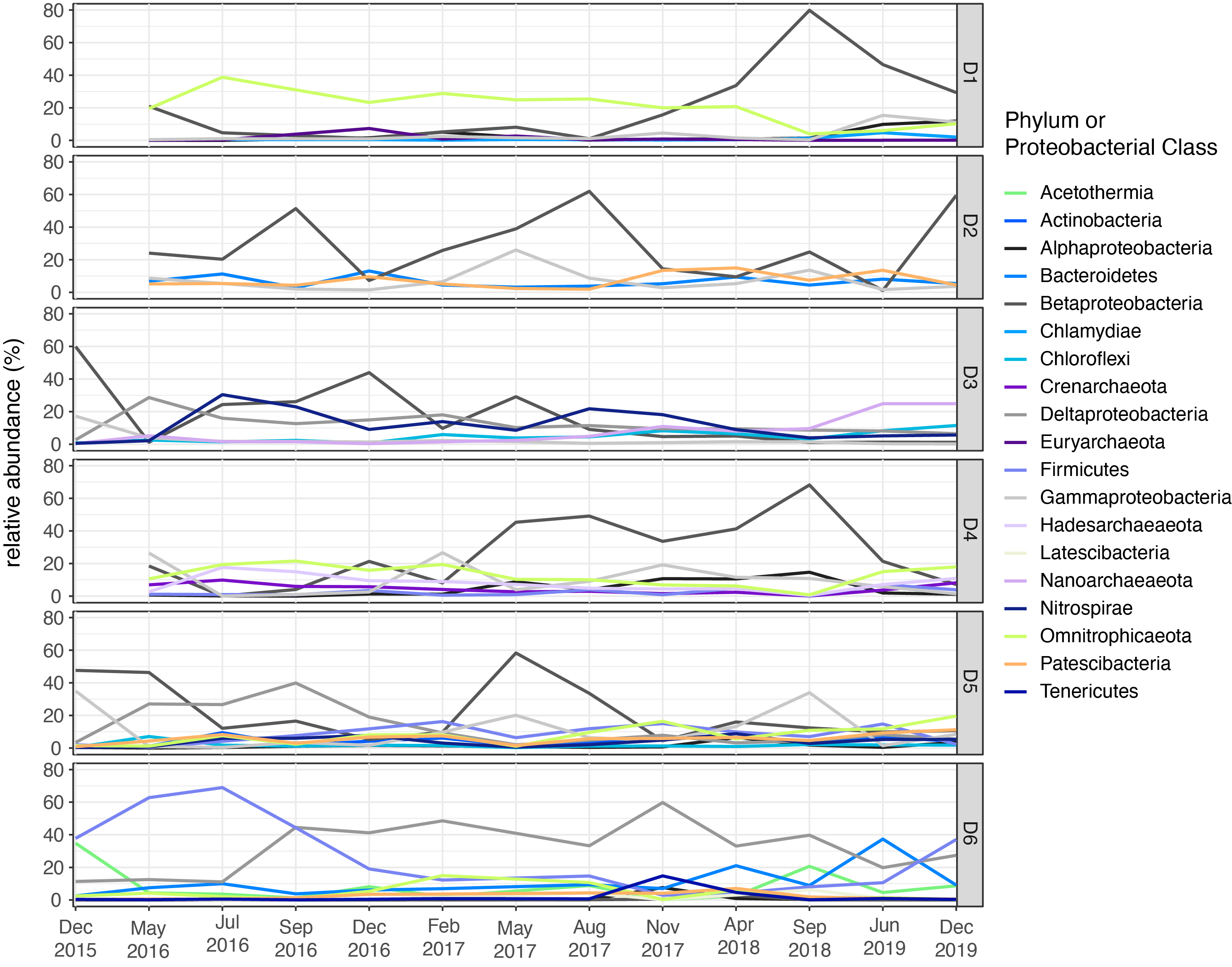

**Supplementary Figure 3:** Line plots of highly variable phyla and Proteobacterial classes over the course of the experiment divided by site. The variability cut off was var:abu > 1.5 and unassigned taxa were excluded. The colors denoting taxa are consistent with those used in Fig. 1.

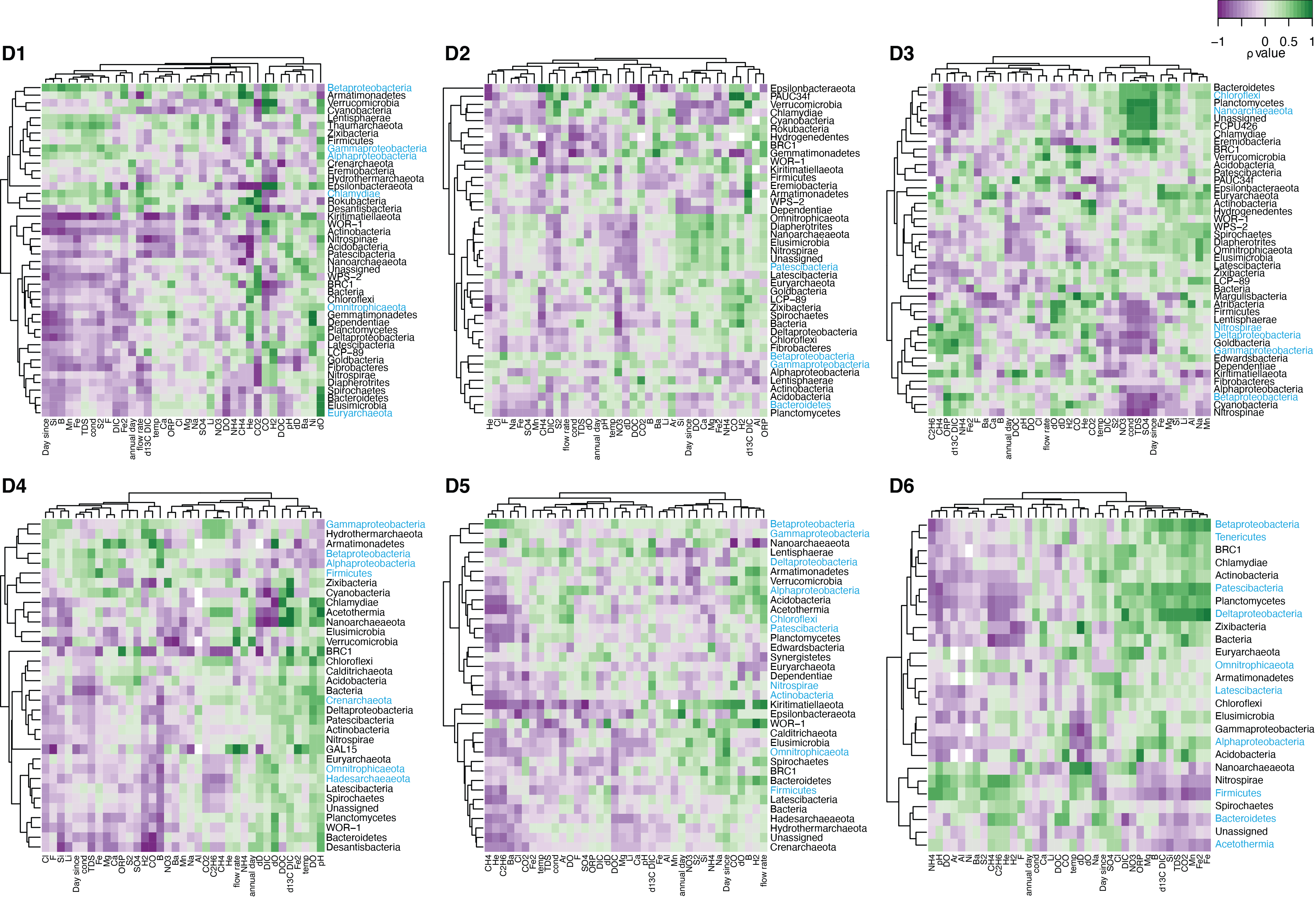

**Supplementary Figure 4:** Heat maps for D1-D6 each depicting a correlation matrix of microbial taxa and geochemical parameters all time points. Taxa which were present in less than 60% of the samples for a given site were excluded. Correlations were produced with the cor() function in R using a pairwise observations and the Spearman’s rank correlation. Hierarchical clustering groups taxa and geochemical parameters with similar behavior. Blue taxa names are highlighting variable taxa for each site shown in Fig. 6 and discussed below.

**Supplementary files**:

HTML version of Sub Fig 1 (average OTU bars)

HTML version of figure 3 (NMDS)

Geochemistry data table (.csv)
